# Genome-wide sequence data show no evidence of admixture and introgression among pollinator wasps associated with a community of Panamanian strangler figs

**DOI:** 10.1101/2020.12.09.418376

**Authors:** Jordan D. Satler, Edward Allen Herre, Tracy A. Heath, Carlos A. Machado, Adalberto Gómez Zúñiga, John D. Nason

## Abstract

Interactions between plants and their animal pollinators can shape processes of divergence and gene flow within associated lineages. For example, in the obligate mutualism between figs (*Ficus*) and fig pollinator wasps (family Agaonidae), each wasp species typically pollinates a single fig species, potentially reinforcing reproductive isolation among different wasp species. Multiple pollinator species, however, can sometimes reproduce in the same host fig species, potentially enabling hybridization and introgression between wasp species. In a community of Panamanian strangler figs (section *Americana*), we use genome-wide ultraconserved element (UCE) loci to estimate phylogenetic relationships and test for hybridization and gene flow among 19 pollinator species associated with 16 host fig species. Previous studies showing ongoing pollinator sharing and a history of pollinator host switching are consistent with documented genetic admixture in their host figs. Here we investigate if host sharing and a dynamic evolutionary history including host switching has also resulted in hybridization and gene flow between pollinator species. Phylogenetic analyses recover strong support for well-delimited wasp species coupled with high interspecific divergence. There is no evidence for ongoing hybridization or introgression, even among pairs of pollinator species currently reproducing within the same host. In contrast to work suggesting admixture among Panamanian host figs, we conclude hybridization and interspecific gene flow have not been important processes shaping the evolutionary history of their pollinating wasps.

## Introduction

Hybridization produces offspring exhibiting novel genetic combinations derived from divergent parental genomes. Introgression results in genetic material moving between species. The influx of genetic material into a lineage introduces new alleles and multilocus combinations that, depending on the situation, can be either harmful (Rhymer and Simberloff, 1996) or beneficial (Dowling and Secor, 1997). In many lineages, strong pre- and postzygotic barriers can limit hybridization and reinforce species boundaries (*e.g.*, Nosil et al., 2005). Evidence from next-generation sequencing, however, has revealed that hybridization and introgression have occurred throughout the evolutionary history of the tree of life (Mallet et al., 2016; Taylor and Larson, 2019). Identifying the ecological processes that either maintain or undermine species boundaries is useful for generating testable hypotheses concerning factors that affect the evolutionary importance of hybridization and introgression (Twyford and Ennos, 2012).

Host specialization in plant-associated insects generally leads to a reduction in interspecific interactions, limiting opportunities for hybridization. This is particularly true for brood pollination mutualisms (*e.g.*, figs and fig wasps, yuccas and yucca moths, globeflowers and globeflower flies) that often exhibit relatively species-specific host–pollinator relationships (Hembry and Althoff, 2016). Figs (*Ficus*; ca. 800 species) and their obligately-associated pollinating wasps (family Agaonidae) present evolutionarily replicated systems that often vary in their potential for hybridization. While most figs appear to be associated with a single, distinct pollinator fig wasp species, there are many cases where fig species are pollinated by multiple, co-occurring pollinator species (*e.g.*, Wiebes, 1995; Kerdelhué et al., 1997; Molbo et al., 2004; Machado et al., 2005; Jackson et al., 2008; Wang et al., 2016; Sutton et al., 2017). Further, at the species range scale about 30% of figs are pollinated by more than one wasp species (Yang et al., 2015). What is certain is that when wasps share the same host—especially when different wasp species reproduce in the same individual figs—there are opportunities for hybridization in either the host fig, the pollinator wasp, or both.

Fig syconia—urn-shaped, enclosed inflorescences—define the genus *Ficus*. Chemical, behavioral, and morphological traits are thought to be important for the female pollinating wasp to identify, locate, and enter a receptive fig (Herre et al., 2008). When a female wasp arrives at a fig tree whose receptivity is signaled by fig floral volatiles (Van Noort et al., 1989; Ware et al., 1993; Grison-Pigé et al., 2002; Hossaert-McKey et al., 2010), she must enter a syconium through a small terminal pore (the ostiole) that excludes other insects. Once inside, the foundress wasp pollinates flowers, oviposits into a subset of them, and dies within the fig (Janzen, 1979). Offspring then develop over several weeks inside the galled, univolulate fig flowers (Galil and Eisikowitch, 1968). Male wasps emerge from their galls, locate galls that contain female wasps, and chew holes in the galls to expose females for mating. After mating, female pollinators emerge from their galls, gather pollen from male flowers in the fig, exit the syconium, and disperse. Female wasps routinely travel many kilometers to encounter another receptive fig inflorescence—on typically a conspecific fig tree—to reproduce (Nason et al., 1998; Ahmed et al., 2009).

Given this reproduction biology, if a fig is visited by a single pollinator wasp (foundress), all pollinator wasp offspring will be her direct descendants—haploid sons and diploid daughters— and mating within the fig will be between siblings. In contrast, if two or more foundresses enter and oviposit within the fig, there are opportunities for non-sibling mating. If these foundresses represent different species, then there is the opportunity for hybridization. Since figs vary substantially in their characteristic foundress numbers (Herre, 1989), opportunities for outcrossing and hybridization in pollinator wasps vary among species. Thus, to potentially have hybridization and interspecific gene flow between pollinators, multiple foundresses must occur within the same fig syconium, and those foundresses must represent different species.

There are approximately 120 described species of strangler figs (*Ficus* subgenus *Urostigma*, section *Americana*) in the Neotropics (Berg, 1989). Figs in this group are pollinated by wasps from the genus *Pegoscapus* (family Agaonidae). Previous studies have emphasized specific associations of particular wasp species with particular host fig species, with its suggestion of strict sense co-speciation of wasp and fig species (*e.g.*, Ramírez, 1970; Bronstein, 1987). Yet, cophylogenetic studies of strangler figs and their pollinators—throughout the Neotropics (*e.g.*, Cruaud et al., 2012) and within Panama (*e.g.*, Jackson et al., 2008)—have identified highly discordant phylogenetic patterns. This indicates that over evolutionary timescales wasps are not particularly host-specific and host switching is an important evolutionary process in this group. In particular, using genome-scale data coupled with a model-based approach, Satler et al. (2019) estimated host switching to be the most important process generating phylogenetic patterns in a community of Panamanian strangler figs and pollinating wasps. Although contemporary associations in this community are predominantly species-specific, phylogenomic data indicate a dynamic evolutionary history punctuated by host-switching events, with these events creating opportunities for hybridization among both associated pollinator and fig species.

Hybrid pollinator wasps have been detected in the Panamanian community of strangler figs. Molbo et al. (2003) sampled emerging pollinators from nine fig species. Using mitochondrial sequence data and microsatellite genotyping, they identified successful F1 hybrids in 4 of 457 (0.9%) broods sampled from one fig species, *Ficus obtusifolia*. These rare F1s occurred between the two frequently co-occurring sister wasp species, *Pegoscapus hoffmeyeri* sp. *A* and *Pegoscapus hoffmeyeri* sp. *B*, that locally pollinate *Ficus obtusifolia*. While occasional wasp hybridization exists, it is unclear whether successful introgression occurs, and raises the question of whether or not hybrids exhibit anything other than negligible fitness. Further, Molbo et al. (2004) recovered two diploid males (out of 18 males sampled from four broods containing hybrid females) among hybrid pollinators associated with *F. obtusifolia*. They suggest a breakdown in the sex determination mechanism caused by hybridization as a factor potentially reducing fitness among hybrid pollinators. Molbo et al. (2004) also collected microsatellite data to test for hybridization between two more distantly related wasp species associated with *Ficus popenoei*. Even though fruits with two foundresses are relatively common for *F. popenoei*, among 255 sampled wasps they recovered no evidence of hybridization. These studies of co-occurring pairs of pollinator species suggest that while hybrids may be occasionally produced, they are functionally sterile and do not contribute genetic material to the next generation.

Here we estimate phylogenetic relationships and test for hybridization and introgression among 19 species of pollinating wasps associated with a community of 16 species of Panamanian strangler figs. First, we use genome-wide sequence data representing ultraconserved element loci to estimate a phylogeny of all sampled fig wasps to test if individual wasps sampled from the same host fig species cluster together in phylogenetic space. Next, focusing on five pairs of wasp species known to pollinate, develop, and mate within figs of the same host fig species, we test for evidence of hybridization and gene flow among these co-occurring pollinators. We then test for hybridization and gene flow more broadly among all pollinator species. Finally, we consider the ecological and evolutionary processes shaping diversification dynamics in these pollinator wasps and how they compare with diversification dynamics in their associated host figs.

## Materials & Methods

### DNA sampling and sequencing

Pollinator wasps were collected from strangler fig species in the vicinity of Barro Colorado Island Nature Monument in the Canal Zone of central Panama (Table 1). Wasps were allowed to emerge from mature figs in the lab and then stored in 95% ethanol or RNALater for DNA extraction and analysis. A single wasp was selected per fig fruit for sequencing to ensure independence among samples. We also included four pollinator wasps from an undescribed species (*Pegoscapus* sp.) associated with the Mexican strangler fig *Ficus petiolaris* to serve as an outgroup; a broader survey has revealed this pollinator to be sister to all other Central American *Pegoscapus* wasps (J. Satler, unpublished data). Including the outgroup, we generated sequence data from 176 wasp samples representing 20 pollinator species, with an average of 8.8 individuals per species (Table 1). Genomic DNA was extracted with a Qiagen DNeasy Kit (Qiagen Inc., Valencia CA, USA). Illumina libraries were generated with a KAPA Hyper Prep kit. Samples were sheared to an average size of ∼450 bp on a Covaris sonicator. Following library construction described in Glenn et al. (2019), we grouped samples in sets of eight, and hybridized biotinylated RNA probes to capture targeted loci. We used the hymenopteran probe v2 set of Branstetter et al. (2017) to target 2590 ultraconserved element (UCE) loci. After probe hybridization and library amplification, size distributions were checked on a Bioanalyzer and libraries were combined in equimolar concentrations for sequencing. Libraries were sequenced on an Illumina sequencer targeting 150 bp paired-end reads.

**Table 1:**
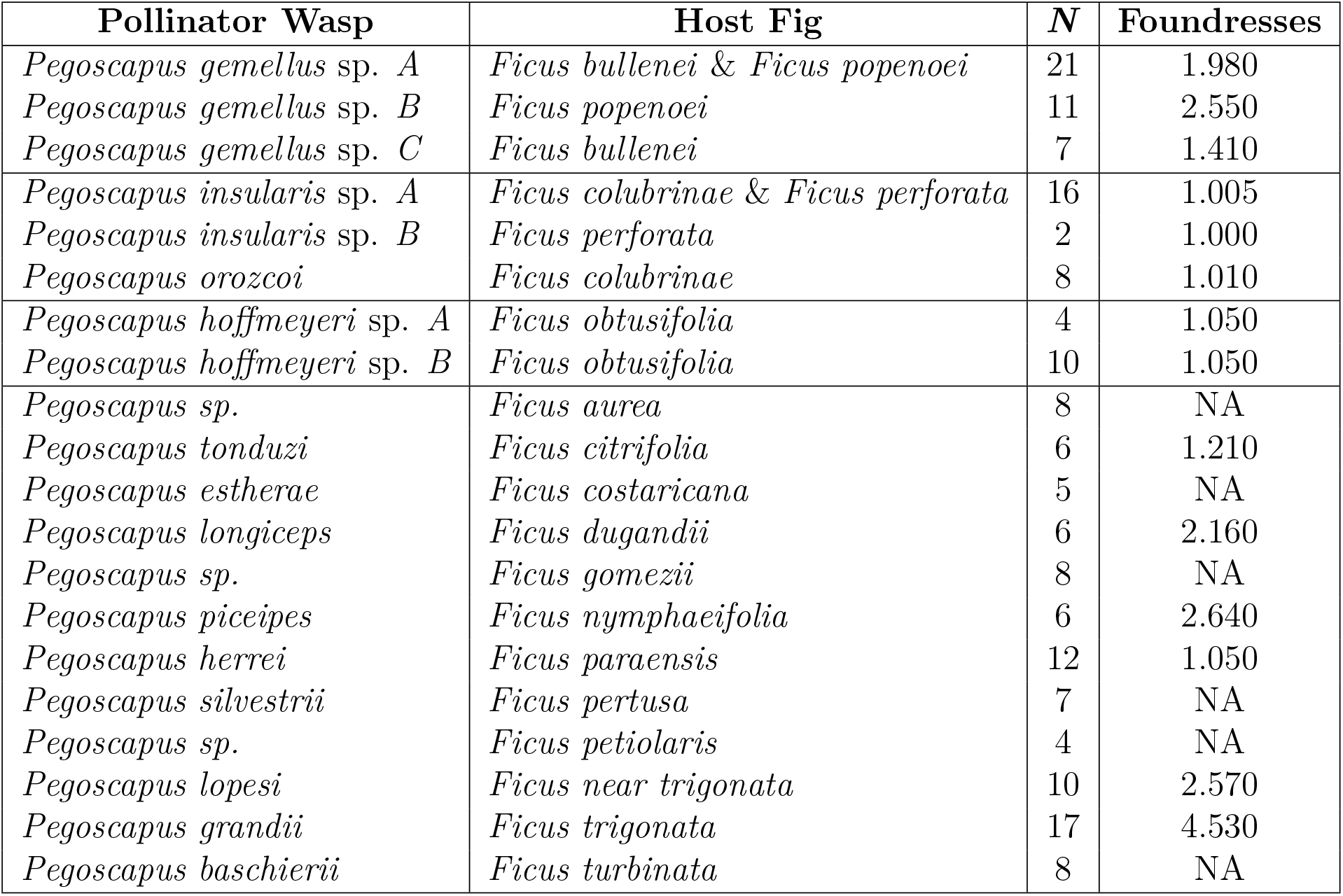
Pollinator wasp sampling. Information includes pollinator species, host fig species, the number of individual wasps sequenced, and the average number of foundresses per host(s). Foundress information, when present, is from Herre (1989). For figs that share a pollinator species, foundress numbers were averaged between the host figs.

### Data processing

We used Phyluce v1.6.7 (Faircloth, 2015) to process raw sequence reads and generate data sets for downstream analysis. Sequence reads were cleaned with illumiprocessor v2.0.9 (Faircloth, 2013), a wrapper around Trimmomatic v0.39 (Bolger et al., 2014). Cleaned reads were assembled into contigs with Trinity v2.0.6 (Grabherr et al., 2011). Contigs were then aligned to the hymenopteran v2 UCE locus set to filter non-specific sequences. Loci were subsequently aligned with MAFFT v7.407 (Katoh and Standley, 2013), and ambiguously aligned sites were removed with Gblocks v0.91b (Castresana, 2000) using default parameters. All loci sampled for a minimum of 70% of individuals were retained in the final data set. Additionally, we generated a data set of phased alleles for certain downstream analyses following the outline provided by Andermann et al. (2018). Taking our aligned loci (before the use of Gblocks), we mapped cleaned sequence reads for each individual to the UCE locus set using BWA-MEM (Li, 2013) in bwa v0.7.17 (Li and Durbin, 2010). We then phased the data with samtools v1.9 (Li et al., 2009) using the phase command, calling two alleles per individual per locus. Loci were then realigned and cleaned as described above, with loci sampled for a minimum of 70% of individuals retained for downstream analysis.

### Phylogenetics

We used both concatenation and coalescent-based species tree methods to estimate phylogenetic relationships among the fig wasps. We first estimated a concatenated phylogeny using maximum likelihood (ML) in IQ-TREE v1.6.12 (Nguyen et al., 2015; Chernomor et al., 2016). This approach allows us to test species monophyly as individuals are treated as tips in the tree. The data set was partitioned by UCE locus, with each partition estimated under the GTR + *γ* substitution model. Nodal support values were generated through 1,000 repetitions of the ultrafast bootstrap approximation (Hoang et al., 2018).

We used two coalescent-based approaches to estimate a species tree. Rather than assuming all genes evolved under the same tree topology, these methods allow for discordance between the gene histories and species tree by explicitly accounting for the biological process of incomplete lineage sorting. First, we used the program SVDquartets (Chifman and Kubatko, 2014) as implemented in PAUP* v4.0a166 (Swofford, 2003). SVDquartets uses site patterns in the nucleotide data to estimate a phylogeny under the multi-species coalescent model. We used SVDquartets in two ways. We initially estimated a lineage tree (SVDQ_*LT*_), where all individuals are represented as tips in the tree, to confirm species assignment and test species monophyly. We then assigned individuals to species *a priori* and estimated a species tree (SVDQ_*ST*_). For both SVDquartets analyses, we evaluated all quartets and used standard bootstrapping to generate nodal support values. Second, we used the program ASTRAL-III v5.6.3 (Zhang et al., 2018) to estimate a species tree. Instead of using site patterns in the nucleotide data, ASTRAL-III uses gene trees as input to estimate a species tree. Maximum likelihood gene trees were first estimated in IQ-TREE. For each locus, IQ-TREE selected the substitution model of best fit with ModelFinder (Kalyaanamoorthy et al., 2017) using Bayesian Information Criteria (BIC). We assigned individuals to species *a priori* (Rabiee et al., 2019), and then used the ML gene trees as input to estimate a species tree. Nodal support values were quantified with local posterior probability values (Sayyari and Mirarab, 2016).

Previous work in the Panamanian system has recovered pollinator species as monophyletic and has not brought into question species validity (Machado et al., 2005; Jackson et al., 2008; Satler et al., 2019). Given the typical pattern of small intraspecific divergence coupled with large interspecific divergence for these wasps, we wanted to ask what proportion of loci recover each species as monophyletic. If the pollinators have been evolving in isolation over evolutionary time, we would expect the species to be monophyletic for all or nearly all sampled loci. In contrast, if interspecific gene flow has been an important process in the evolution of this system, we would expect interacting species to show a lack of monophyly for numerous loci. We used DendroPy v4.4.0 (Sukumaran and Holder, 2010) to count the proportion of gene trees (as estimated above in IQ-TREE) for which a species was monophyletic. We required that at least two individuals were sequenced for a species for a given gene tree to assess monophyly.

### Population genetics

In addition to estimating phylogenetic relationships and testing for monophyly, we were interested in understanding the population genetics of the pollinator species. In particular, we wanted to test if genetic diversity within a species varied with average foundress number. If outbreeding is common among species that contain multiple foundresses per fruit, we would expect to see a positive correlation between genetic diversity and average foundress number. We used DendroPy to calculate nucleotide diversity (*π*), number of segregating sites, and Watterson’s theta (per site) for all pollinator species. For species with foundress number data, we used Pearson’s correlation test in R v3.5.2 (R Core Team, 2018) to test for a correlation between average number of foundresses and summary statistic.

### Testing for monophyly with mitochondrial DNA

To complement analyses and inference with the genome-wide data from the nuclear genome, we used mitochondrial DNA (mtDNA) to test if pollinator species were monophyletic in the mitochondrial genome. Even when not targeted, mitochondrial DNA is often collected using sequence capture approaches (do Amaral et al., 2015; Barrow et al., 2017). We used NOVOPlasty v3.8.3 (Dierckxsens et al., 2017) to identify mitochondrial reads and generate haplotypes from the sequencing files. Since there is no *Pegoscapus* mitochondrial reference genome available, we aligned reads to a *Pegoscapus* cytochrome oxidase I (COI) mtDNA sequence (AY148119). We used default settings with an initial kmer value of 39. For samples that either did not generate any matches or produced obvious sequencing errors, we lowered the kmer value to 23. Using a lower kmer value can help with low coverage data, appropriate here since we were only targeting regions of the nuclear genome. Finally, for samples that did not recover any mtDNA, we aligned reads to a longer *Pegoscapus* COI sequence of a different species (JN103329) using default settings. Mitochondrial haplotypes were aligned with MAFFT and edge trimmed to match the primary seed sequence to reduce missing data. We estimated a maximum likelihood gene tree in IQ-TREE, using ModelFinder to select the substitution model and 1,000 repetitions of the ultrafast bootstrap approximation to generate nodal support. If patterns of monophyly differ between the nuclear and mitochondrial genomes, the cytonuclear discordance would suggest introgression or processes other than genetic drift generating these discordant patterns.

### Testing for admixture and gene flow

Detailed studies of Neotropical strangler figs and their pollinators have revealed exceptions to the typical 1:1 association between fig and wasp species (Molbo et al., 2003, 2004; Machado et al., 2005). Cases of host sharing, where multiple pollinator species are associated with the same host, are of particular interest because they present clear opportunities for hybridization and introgression. Five cases of host sharing have been described in central Panama: (i and ii) *Ficus bullenei* and *Ficus popenoei* share a pollinator, *Pegoscapus gemellus* sp. *A*, which co-occurs with *Pegoscapus gemellus* sp. *C* in *F. bullenei* and *Pegoscapus gemellus* sp. *B* in *F. popenoei*, resulting in two hosts each with two co-occurring pollinators, (iii and iv) *Ficus colubrinae* and *Ficus perforata* share a pollinator, *Pegoscapus insularis* sp. *A*, which co-occurs with *Pegoscapus orozcoi* in *F. colubrinae* and *Pegoscapus insularis* sp *B* in *F. perforata*, again resulting in two hosts each with two co-occurring pollinators, and (v) *Ficus obtusifolia* is pollinated by two co-occurring species, *Pegoscapus hoffmeyeri* sp. *A* and *Pegoscapus hoffmeyeri* sp. *B*. We refer to these three systems as BP (i and ii), CP (iii and iv), and O (v) (see Figure 1). They include two pollinator species associated with two hosts each, and six pollinator species that are host specific. It is the interaction of wasps within the same fig hosts that provide the greatest opportunities for hybridization and gene flow.

**Figure 1:**
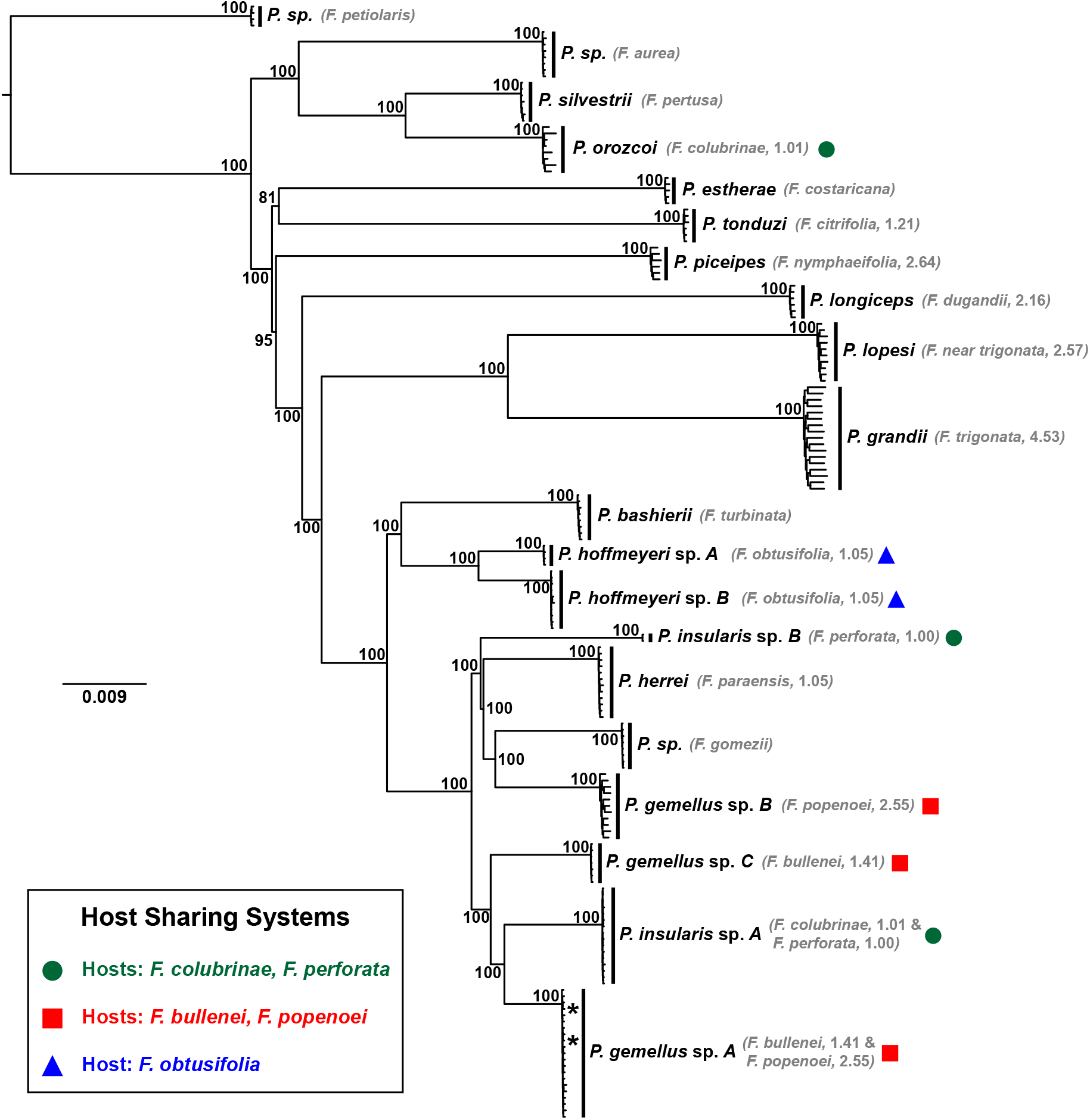
Maximum likelihood phylogeny representing relationships among *Pegoscapus* wasps. Host fig species are displayed next to their associated wasp species, with average foundress number data included when available. See Herre (1989) for details. Node labels represent bootstrap support values. The insert shows the three systems with co-occurring pollinators, denoted by a circle (green), square (red), or triangle (blue). Two individuals (represented by asterisks) of *Pegoscapus gemellus* sp. *A* (hosts: *Ficus bullenei*/*Ficus popenoei*) were sampled from *Ficus dugandii*, which is not its normal host. The undescribed pollinator associated with *Ficus petiolaris* was used to root the tree.

The fig species with co-occurring pollinators vary in their average number of foundress wasps per fig (Herre, 1989). Variation in foundress number enables predictions about the pollinator taxa most likely to contain histories of hybridization and gene flow. *Ficus colubrinae* and *F. perforata* are nearly always single foundress (Table 1). Despite two pollinator species associated with each fig species, we expect little opportunity for hybridization since more than one foundress rarely occupies the same syconium. *Ficus obtusifolia* is mostly single foundress, averaging 1.21 foundresses per fruit, although 16% of sampled fruits had between two and four foundresses (Herre, 1989). We expect intermediate opportunities for hybridization between the two pollinators associated with this host. Finally, *F. bullenei* averages 1.41 foundresses per fruit and *F. popenoei* averages 2.55 foundresses per fruit, providing frequent opportunities for pollinator co-occurrence and, potentially, hybridization between their two pairs of wasp species. Together, the five pairs of host-sharing pollinators vary substantially in their expected opportunities for hybridization and interspecific gene flow.

We used two approaches to test for hybridization and admixture between co-occurring pollinator species. First, we used principal components analysis (PCA). If sampled individuals are recent hybrids or backcrosses to parental lineages, we would expect to see individuals intermediate between clusters representing parental lineages. Further, if introgression has been extensive, there should be little separation of clusters in PCA space. PCA analysis was conducted with the dudi.pca function in the R package ade4 (Dray et al., 2007). Missing data were replaced with mean values of allele frequencies within a given species. We plotted the first two axes of variation to visualize genetic clustering of individuals. Second, we used Structure v2.3.4 (Pritchard et al., 2000) to cluster individuals into genetic groupings. Structure clusters individuals by maximizing Hardy-Weinberg equilibrium within clusters and minimizing Hardy-Weinberg equilibrium among clusters. Evidence for recent hybridization and introgression would be reflected by individuals containing multilocus genotypes sampled from multiple clusters. Our choice of the number of clusters (*K*) was informed by previous work identifying the wasp species within these systems (Molbo et al., 2003; Jackson et al., 2008). For *F. bullenei*/*F. popenoei* (BP) and *F. colubrinae*/*F. perforata* (CP), we used a *K* value of 3; for *F. obtusifolia* (O), we used a *K* value of 2. We used the admixture model, and allowed allele frequencies to be correlated among populations. Each analysis had a burnin of 100,000 steps, followed by 500,000 MCMC reps, and was replicated 10 times. Results were processed with the R package pophelper v2.2.7 (Francis, 2017).

For the pollinators in each host system (*F. colubrinae* and *F. perforata*, *F. bullenei* and *F. popenoei*, *F. obtusifolia*), genetic data were processed independently to generate phased data sets. By processing each taxon set independently, we retained loci with a minimum of 70% for the taxa of interest only. We followed the same procedure as outlined above for generating the phased data sets, and used unlinked SNPs for both the PCA and Structure analyses. To generate these data sets, we first scanned our phased aligned taxon-specific UCE loci for variable sites. Within a locus, we then selected the SNP that had the highest sample coverage. If a locus had multiple SNPs with equal sample coverage, one of those SNPs was selected at random.

Tests of hybridization among co-pollinators provide biologically motivated hypotheses where opportunities for interspecific mating are most likely. If gene flow among these species has been recent or ongoing, we would expect to detect that signal and infer a history of hybridization. This approach, however, is limited to specific sets of taxa based on present day associations. Given the history of host switching in this system, and accounting for a potential dynamic process of host–pollinator associations through time, it would be useful to test for hybridization among all sampled taxa.

To test if historical introgression has been an important process in this community of pollinators, we used TreeMix v1.13 (Pickrell and Pritchard, 2012). TreeMix is a maximum likelihood approach that uses allele frequency data to construct a population graph and places hybridization events on that graph for populations with the least fit to a tree model. Since the number of hybridization events are specified *a priori*, we can test models varying the number of hybridization events to determine a model of best fit. We used the total variance explained by the model to inform the number of hybridization events that provides a best fit for this community of pollinator species. We used allele frequency data representing unlinked, biallelic SNPs from the phased data set from all pollinator species. To remove effects of missing data, we first identified biallelic SNPs within each locus that were sampled for at least one individual per species. For loci with multiple SNPs, we selected a single SNP with the highest coverage. If a locus had multiple SNPs with equal sample coverage, one SNP was selected at random. We then estimated a maximum likelihood population graph in TreeMix, allowing between zero and five migration events.

Although additional approaches are often used to test for hybridization and introgression, they were not applicable for our data set. For example, *D*-statistics (Green et al., 2010; Durand et al., 2011), often referred to as ABBA/BABA tests, have been widely used to infer a signal of introgression among numerous clades across the tree of life (*e.g.*, Eaton and Ree, 2013; Streicher et al., 2014; Meier et al., 2017; Hughes et al., 2020; Pulido-Santacruz et al., 2020). *D*-statistics, however, can not estimate gene flow between sister species, and our biologically informed hypotheses do not lend themselves well to the strict phylogenetic pattern (((P1,P2),P3),OG) typically required for testing if one species (P3) has hybridized with another (either species P1 or species P2). In addition, if we use an agnostic approach and test all possible combinations of taxa, that would require 969 tests (19 choose 3), and would potentially lead to artifacts from unsampled (“ghost”) lineages given the subsampling approach (Pease and Hahn, 2015).

## Results

### DNA sampling and sequencing

We generated 468,733,333 total sequence reads from the 176 fig wasp samples, resulting in an average of 2,663,258 (±1, 273, 010) reads per individual. Assembly in Trinity resulted in an average of 139,295 (±80, 671) contigs per individual. After filtering and retaining contigs that matched UCEs in the probe set, we retained an average of 1,590 (±179) UCE loci per individual. Filtering loci for those that contained at least 70% taxon sampling, we retained 1,504 loci for the complete taxonomic data set. In the three systems (processed independently) with shared and unique pollinators, at 70% sequencing threshold, we retained 1,438 (BP), 1,388 (CP), and 1,390 (O) loci for analysis.

### Phylogenetics

When individuals are treated as tips in the tree, phylogenetic analyses recover each pollinator species as monophyletic with strong support. This result is consistent with both concatenation (Figure 1) and in the coalescent-based lineage tree analysis with SVDquartets (SVDQ_*LT*_, Figure S1). To further investigate, we tested species monophyly for each sampled UCE locus. All species were monophyletic for either all or nearly all loci (Table 2). 58% of species were monophyletic in at least 95% of loci, while 84% of species were monophyletic in at least 90% of loci. Only three species were monophyletic at less than 90% of their loci, with the lowest proportion being 82% (*P. gemellus* sp. *A* (hosts: *F. bullenei*/*F. popenoei*). For species with foundress data (see Table 1), there is no significant correlation between average foundress number and proportion of monophyletic loci (Pearson’s correlation test, *r* = 0.212, *P* = 0.466). Although we do not distinguish the causes of non-monophyly (*e.g.*, deep coalescence, introgression, gene tree estimation error), these conservative estimates are considerably high and provide strong support for the validity of the wasp species and their isolation over evolutionary time.

**Table 2:**
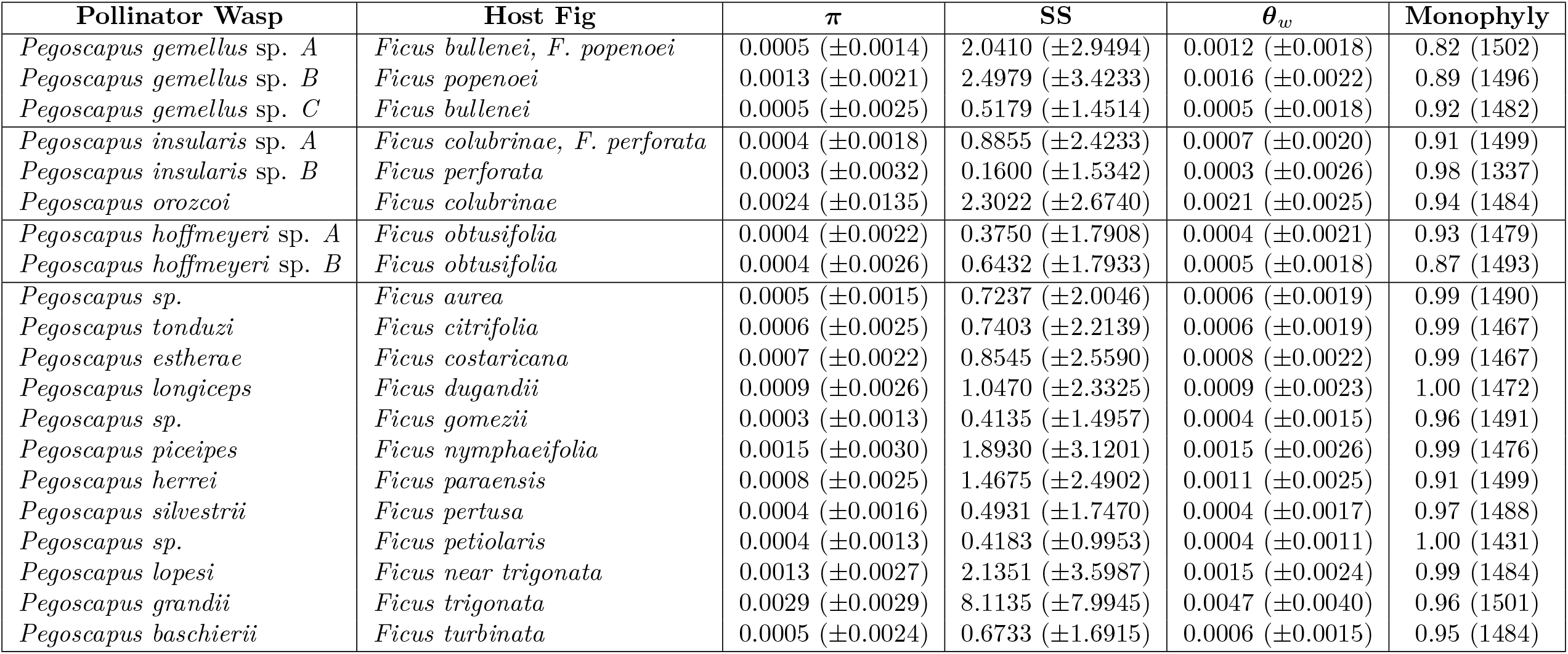
Population genetic summary statistics for the pollinator wasp species. Summary statistics include nucleotide diversity (*π*), number of segregating sites (SS), and Watterson’s theta per site (*θ*_*w*_). Monophyly shows the proportion of gene trees for which a pollinator species is monophyletic. A gene tree was tested for monophyly when two or more sequences for a given species were sampled, with total trees tested per species in parentheses.

Phylogenetic relationships are congruent across all three methods. For the concatenated tree, all nodes are strongly supported (most have a bootstrap value of 100), with a general pattern of small intraspecific divergence and large interspecific divergence (Figure 1). Two *P. gemellus* sp. *A* (hosts: *F. bullenei*/*F. popenoei*) individuals were sampled from *Ficus dugandii*, indicating an example of non-host specificity. In the species tree analyses SVDQ_*ST*_ (Figure 2) and ASTRAL-III (Figure S2), there is strong support for most relationships among species, although a few nodes in both trees are weakly supported. The minor differences in the phylogenetic analyses occur at nodes with low support or at short internal branches. For example, IQ-TREE supports *Pegoscapus herrei* (host: *Ficus paraensis*) sister to (*Pegoscapus sp.* (host: *Ficus gomezii*), *P. gemellus* sp. *B* (host: *F. popenoei*)), with *P. insularis* sp. *B* (host: *F. perforata*) sister to this clade, while SVDQ_*ST*_ and ASTRAL-III places *P. herrei* (host: *F. paraensis*) as sister to a clade of the other three species. In addition, IQ-TREE and SVDQ_*ST*_ place *Pegoscapus tonduzi* (host: *Ficus citrifolia*) sister to *Pegoscapus estherae* (host: *Ficus costaricana*) with weak support, while ASTRAL-III places *P. estherae* (host: *F. costaricana* sister to a large clade of fig wasps, with *P. tonduzi* (host: *F. citrifolia*) sister to them.

**Figure 2:**
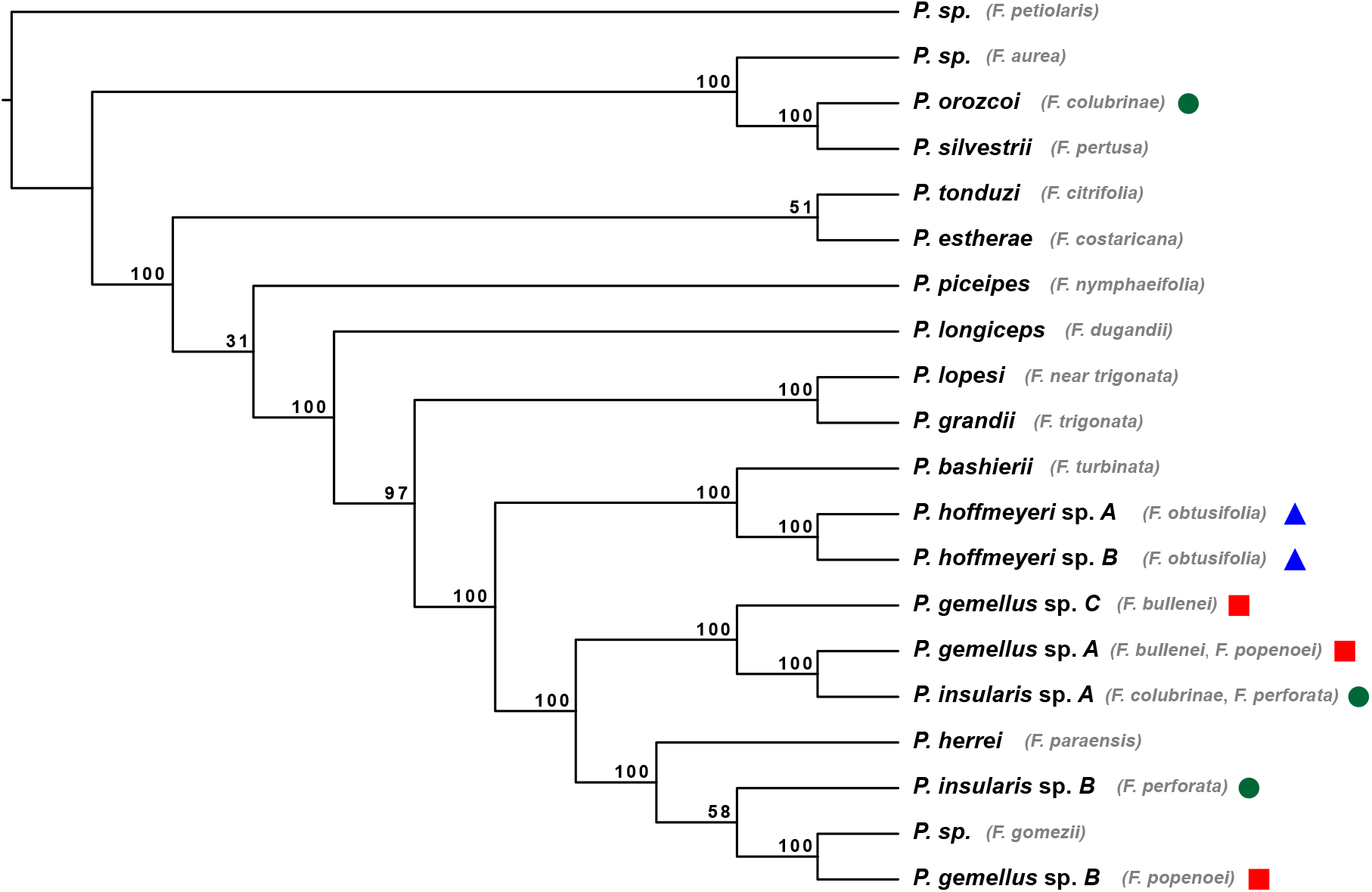
Species tree analysis with SVDquartets (SVDQ_*ST*_). Nodal support is denoted with bootstrap support values. Host fig species are displayed next to their respective wasp species. Co-occurring pollinators are denoted by a circle (green), square (red), or triangle (blue). The undescribed pollinator associated with *Ficus petiolaris* was used to root the tree.

Phylogenetic relatedness varies for the three systems containing co-occurring (*i.e.*, shared and host-specific) pollinators. The two pollinators (*P. hoffmeyeri* sp. *A* and *P. hoffmeyeri* sp. *B*) associated with *F. obtusifolia* are sister taxa in the phylogenies, while the three pollinator species associated with *F. colubrinae* and *F. perforata* are phylogenetically distantly related (Figures 1–2, S1–S2). Pollinator species associated with *F. bullenei* and *F. popenoei* show phylogenetic relatedness intermediate to these two systems, where they are less divergent from one another than the pollinators associated with *F. colubrinae* and *F. perforata* yet are not sister taxa within the tree. Results for these phylogenetic patterns are consistent with both concatenation (Figure 1) and species tree approaches (Figures 2, S2), and highlight a range of evolutionary relatedness among the systems containing co-occurring pollinator species.

### Population genetics

Population genetic statistics were variable across pollinator species, with generally wide ranges and large variances (Table 2). There is a significant positive correlation between average number of foundress per species and genetic diversity (Figure 3). Pearson’s corre

**Figure 3:**
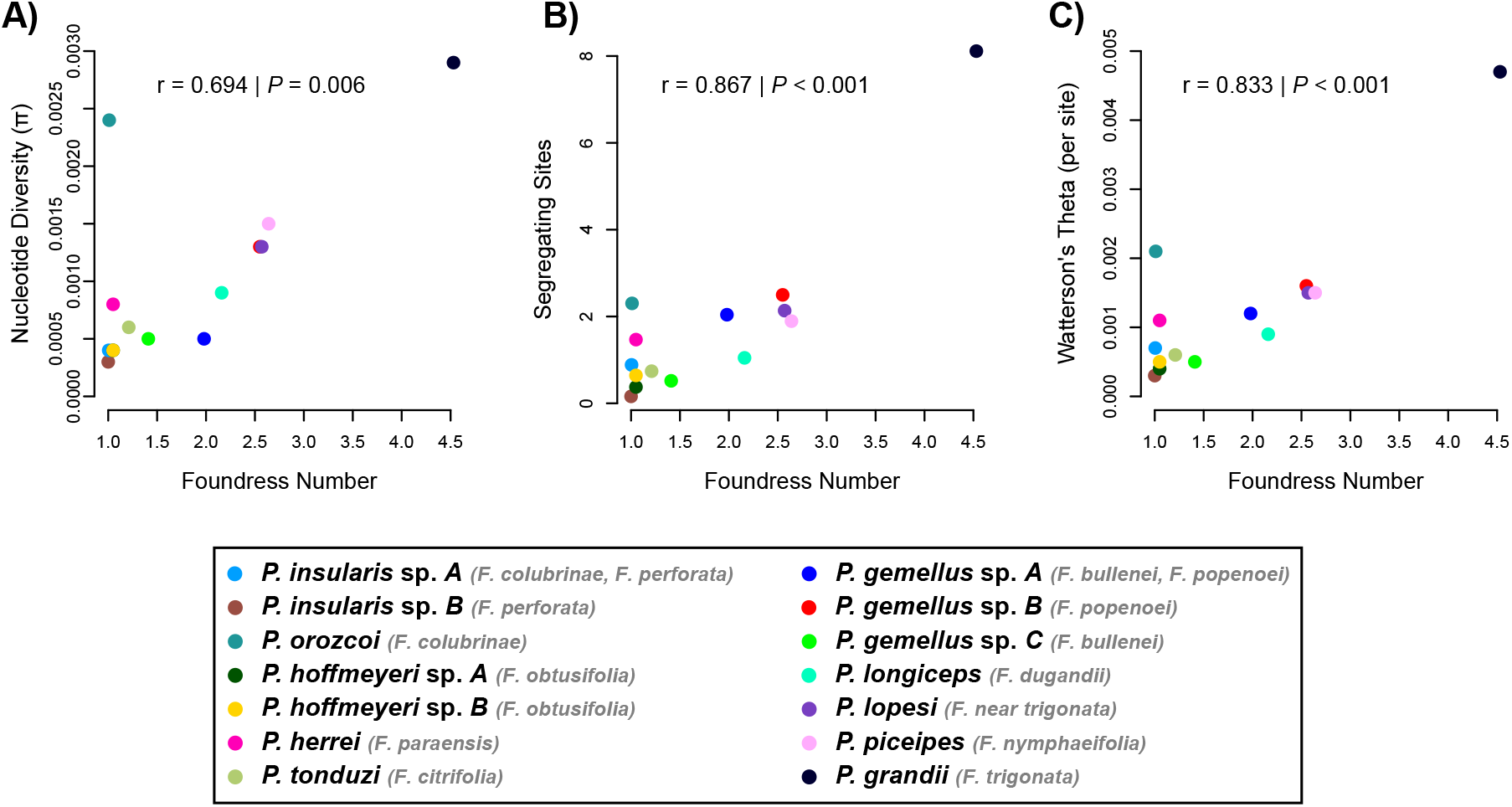
Correlation between population genetic summary statistics and average foundress number per pollinator species. Summary statistics include (A) nucleotide diversity, (B) number of segregating sites, and (C) Watterson’s theta (per site). The Pearson’s product-moment correlation was used to test significance. Fourteen wasp species were included for which we had average number of foundress information (see Herre, 1989). If a wasp had multiple hosts, we used the average for the number of foundresses from the shared hosts. lation test recovered a statistically significant positive correlation for nucleotide diversity (*r* = 0.694, *P* = 0.006), number of segregating sites (*r* = 0.867, *P* < 0.001), and Watterson’s theta (*r* = 0.833, *P* < 0.001) versus number of foundresses.

### Testing for monophyly with mitochondrial DNA

We were able to capture mitochondrial DNA of the COI region from 164 individuals (out of 176). Following alignment and edge trimming, these data comprised 816 base pairs of the COI mtDNA gene. All wasp species are recovered as monophyletic with strong support, with all but one species having a bootstrap value of 100 (Figure S3). There is, however, little support for phylogenetic relationships among species, as most nodal support values are low. Consistent with how individuals cluster within species with the nuclear data, the same clustering of individuals within species is recovered with data from the mitochondrial genome, providing strong support for what we consider a pollinator species in this community. Given the low support values for interspecific relationships, we make no comparisons between the structure of the nuclear genome phylogeny and the mitochondrial gene tree. Comparing how individuals cluster in the nuclear phylogeny and mitochondrial gene tree, however, these data provide no evidence of recent cytonuclear discordance.

### Testing for admixture and gene flow

There was no evidence of admixture or introgression in the three systems (BP, CP, O) containing co-occurring pollinator species. Distinct clusters corresponding to species are recovered in the PCA analyses, with no signal for hybridization (Figure 4). For both BP and CP, wasp species plot in distinct PCA space with little intraspecific variance in comparison to the much larger interspecific variance. The two wasp species in O are well differentiated in PCA axis 1, but show some intraspecific spread in PCA axis 2, although this axis only comprises 0.35% of the variance. These results were further supported in Structure, as individuals were assigned to their respective species with no evidence of admixture (Figure 4). In contrast to seeing shared ancestry indicative of admixture and introgression, each individual maps unambiguously to its respective species. Thus, in systems where opportunity for hybridization is present, we detect none.

**Figure 4:**
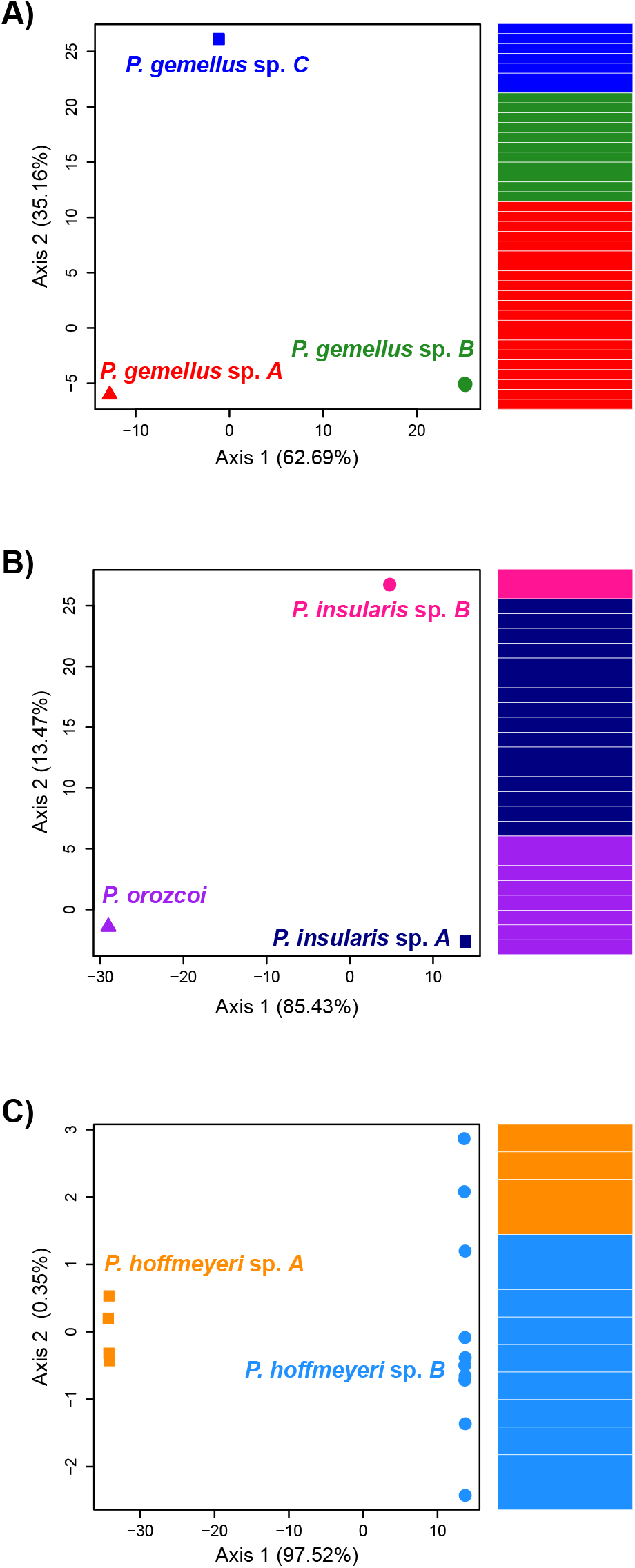
Principal components analysis (PCA) and Structure plots for the three systems where multiple pollinators interact with one or two fig species. Panel A represents three pollinator species interacting with *Ficus bullenei* and *Ficus popenoei*. Panel B represents three pollinator species interacting with *Ficus colubrinae* and *Ficus perforata*. Panel C represents two pollinator species interacting with *Ficus obtusifolia*

TreeMix estimated a population graph consistent with the other phylogenetic approaches (Figure S4). Some differences were seen, mainly at places with short internal branches and low support along the backbone of the tree. Adding migration events to the tree only incrementally improved the proportion of variance explained by the model (Table 3). For example, the population graph with no admixture events explained 98.45% of the variation. Adding one hybridization event to the graph increased the proportion of variation explained by the model to 98.62%. The minimal improvement in models including migration events suggests hybridization is not an important process in this pollinator community.

**Table 3:** Proportion of variation explained by the different models in TreeMix.

| Model | Admixture Events | Percent Variation |
| --- | --- | --- |
| m0 | 0 | 98.45% |
| m1 | 1 | 98.62% |
| m2 | 2 | 98.76% |
| m3 | 3 | 98.81% |
| m4 | 4 | 98.90% |
| m5 | 5 | 98.96% |

## Discussion

We collected genome-wide ultraconserved element (UCE) loci from 19 pollinator wasp species associated with all 16 host species in a strangler fig community from the vicinity of the Panama Canal. We used these data to estimate phylogenetic relationships and test for hybridization and introgression among the pollinators. This group of congeneric pollinators (*Pegoscapus*) has exhibited frequent host switching throughout their shared evolutionary history with their associated fig hosts (Satler et al., 2019), providing opportunities for hybridization in both host fig and pollinator wasp species. Further, rare F1 pollinator hybrids have been detected in this community (Molbo et al., 2003, 2004). We found that all 19 pollinator species are well-delimited genetically and show high interspecific divergence. Even among co-occurring pollinator species in which heterospecific contact is most likely (reproduction within the same fruits of the same host fig species), we detected no evidence for hybridization that led to successful introgression. Our results suggest that successful hybridization and gene flow across species does not contribute meaningfully to the evolutionary history of these Panamanian pollinators, and plays a negligible role in their co-evolution with their host figs.

Because the reproduction of both host and pollinator occur in the same general structure (the fig inflorescence = syconium = fig), there is considerable overlap in the conditions that facilitate hybridization in both partners in this mutualism. In both the host fig and the pollinator wasp, opportunities for hybridization and introgression depend on physical access of heterospecific gametes (*e.g.*, pollen and ovaries for the figs, sperm and eggs for the wasps). The conditions for successful hybridization, however, are less restrictive in the host figs. Here, although foundress wasps must be attracted to the floral volatiles of a different host species and enter and pollinate receptive fig inflorescences, only a single wasp from a heterospecific host need enter a fig to unite heterospecific pollen and ovules. The resulting hybrid seeds must be viable, and introgression is only possible if the hybrid seeds produce viable seedlings that survive to reproduce and backcross, most likely with one of the parental species. The conditions for wasp hybridization are more stringent. Here, two or more foundress wasps from different species must enter the same individual receptive fig inflorescence, with the opportunity for foundress co-occurrence greatly restricted in host species typically producing single-foundress figs (Table 1). After pollination and oviposition, heterospecific male and female offspring must mate, the mated females must successfully disperse to a new receptive fig, and the resulting hybrid offspring must survive, develop, disperse, and reproduce. Introgression requires that these hybrid wasps contribute to successful backcrosses that also must take place within the same fig where other foundresses have successfully oviposited, which will again be restricted in host species with single-foundress figs.

In contrast to our findings in the Panamanian fig wasps, hybridization followed by introgression is strongly suggested in their Panamanian fig hosts (Machado et al., 2005; Jackson et al., 2008). For example, analyses that account for shared ancestral polymorphism show that an isolation-only model provides a poor fit to the data from three Panamanian fig species (Machado et al., 2005). Although based on limited sequence data (three nuclear loci), these authors suggest this poor fit is best explained by hybridization and introgression among the Panamanian figs. Further, Jackson et al. (2008) presented detailed haplotype analyses that were most consistent with widespread hybridization and subsequent introgression across most of the Panamanian figs they studied, which included 14 of the 16 host fig species represented in our study. These findings are consistent with expectations based on the demonstration of pollinator sharing and an evolutionary history of host switching by the pollinators associated the Panamanian figs (Molbo et al., 2003, 2004; Satler et al., 2019). Further, both direct and indirect evidence suggest that hybridization occurs in figs more generally, and that introgression is a potentially important process in the evolutionary history of *Ficus* (Compton, 1990; Machado et al., 2005; Jackson et al., 2008; Compton et al., 2009; Renoult et al., 2009; Cornille et al., 2012; Van Noort et al., 2013; Bruun-Lund et al., 2017). Specifically, cytonuclear discordance between plastid and nuclear genomes suggest ancient hybridization and introgression have played an important role in shaping diversification patterns in *Ficus* (Renoult et al., 2009; Bruun-Lund et al., 2017). Accumulated evidence suggests that host switching and mixed plant–pollinator associations affect the likelihood of hybridization and introgression in each mutualist lineage differently.

Among the pairs of Panamanian wasp species that share the same host fig species, there appears to be no clear pattern to their degree of phylogenetic relatedness. Only *P. hoffmeyeri* sp. *A* and sp. *B* associated with *F. obtusifolia* are recovered as sister species in this community (Figures 1–2, S1–S2), and these are the only two species with evidence supporting rare F1 hybrid events (Molbo et al., 2003, 2004). The other co-occurring pollinators span the phylogenetic breadth of our sampled community, with wasps associated with *F. bullenei* and *F. popenoei* being relatively closely related (but not sister taxa) while wasps associated with *F. colubrinae* and *F. perforata* are distantly related (Figures 1–2, S1–S2). The Panamanian observations are consistent with previously reported estimates that 32.1% of co-occurring pollinators of monoecious fig species are sister species (Yang et al., 2015). Importantly, despite the evidence for rare F1 hybridization events between the two *F. obtusifolia* pollinators (Molbo et al., 2003, 2004), we found no evidence for any backcrosses or introgression in any of the species we studied. Even if our current sampling missed F1 hybrid individuals, if hybridization is important then we would expect to have detected some backcross genotypes or genetic signal of introgression. Given our observations, how might different processes that affect prezygotic (*e.g.*, host recognition, likelihood of sharing individual fig inflorescences) and postzygotic (*e.g.*, developmental incompatibility with the host fig, reproductive incompatibility between wasp species) barriers limit hybridization and introgression among pollinator species?

Host choice by the pollinator fundamentally influences the potential for hybridization and gene flow for both the fig and the wasp. Receptive fig inflorescences produce complex volatile chemical blends recognized by potential pollinators. Generally, the blends produced by different figs are sufficiently distinct for wasps to distinguish among potential hosts (Van Noort et al., 1989; Ware et al., 1993; Grison-Pigé et al., 2002; Hossaert-McKey et al., 2010; Cornille et al., 2012). The degree of affinity for particular wasp species towards particular host blends appears to be an important prezygotic barrier limiting encounter rates of heterospecific wasps. In 17 of the 19 wasp species sampled in the Panamanian community, wasps are predominately attracted to a single host fig species, suggesting high host specificity likely based in large part on recognition of host volatile chemical signals (Table 1) (Molbo et al., 2003, Oldenbeuvingen, Florez, and Herre, unpublished data). Wasp species that share fig hosts, however, are often not sister species, both in this community and more broadly across *Ficus* (Yang et al., 2015). The observation of multiple, phylogenetically divergent pollinator species attracted to the same host fig raises important questions: What processes have shaped the genetics and biochemical pathways of host chemical signal production, of wasp chemosensory detection, and of wasp volatile preference? Addressing this complex series of questions will require studies that explicitly connect host volatile chemical composition with wasp preferences (*e.g.*, antennogram and gene expression studies) within the contexts of detailed phylogenetic and ecological studies, preferably across multiple sites and fig–wasp taxa.

Individual fig fruit crops can exhibit tens of thousands of figs, and effective population sizes of figs consist of hundreds of individuals (Nason et al., 1998; Korine et al., 2000). Even if heterospecific wasps recognize the same host fig species, a precondition for hybridization is that two (or more) heterospecific foundress wasps oviposit successfully in the same individual fig. In the Panamanian strangler figs we studied, the average foundress numbers per fruit ranges between one to four and a half (Herre, 1989). For species pollinated almost exclusively by a single foundress (*e.g.*, Panamanian populations of *F. perforata* and *F. colubrinae*, Table 1), the opportunities for wasp hybridization are limited. In contrast, in fig species commonly having multiple foundresses per fruit, the offspring of pollinators have greater opportunities to encounter heterospecifics in the same fig. Higher mean foundress numbers will also determine wasp population structure which in turn will affect many aspects of these wasp species (*e.g.*, sexual competition and sex ratios, heterozygosity) (Herre, 1985, 1989, 1993; Molbo et al., 2003, 2004). In particular, we found a significant positive correlation between the average number of foundresses and genetic diversity of UCE loci across pollinator species (Figure 3). As was the case with species that show occasional F1 hybrids, however, we detected no evidence of hybridization and introgression in pollinator species that exhibited higher average foundress numbers, including *Ficus nymphaefolia* (2.6 foundresses) and *Ficus trigonata* (4.5 foundresses). Thus, when multiple foundresses are present, outbreeding occurs within species, but outbreeding—and subsequent introgression—does not occur between species.

Postzygotic barriers to successful wasp hybridization include developmental incompatibility with the host fig during the process of gall formation and the development of wasp offspring (Compton, 1990; Compton et al., 2009; Van Noort et al., 2013; Martinson et al., 2014). For example, Moe and Weiblen (2012) experimentally introduced pollinator wasps into figs of a species that are not their normal hosts. Although individuals of the non-normal wasp species were able to oviposit and induce gall development, the galls and the wasp offspring were unable to develop to maturity, suggesting selective constraints on heterospecific wasp development. It appears that when heterospecific pollinator mating does occur, the hybrid offspring exhibit moderately to extremely reduced fitness compared to non-hybrid individuals (Molbo et al., 2003, 2004).

Another potential postzygotic barrier to hybridization and introgression in the wasps is *Wolbachia*. These maternally inherited cytoplasmic bacteria are known to cause drastic reductions of hybrid formation in *Nasonia*, another chalcidoid wasp (Bordenstein et al., 2001). *Wolbachia* are commonly found across insect species (Werren, 1997), and are particularly common in fig wasps (*e.g.*, Haine and Cook, 2005; Sun et al., 2011). Pollinator and non-pollinator wasps associated with the fig community in central Panama show *Wolbachia* occurrence in 59% of wasp species (Shoemaker et al., 2002), including many of the species sampled in this study. In all fig species in which different wasp pollinators co-occur, at least one of the pollinator species shows partial or complete *Wolbachia* infection (Shoemaker et al., 2002), raising the possibility that these bacteria influence the likelihood of successful hybridization.

Many studies of the associations of pollinating wasps with their host figs tend to focus on one or a few species interactions at one or a few locales. The coupling of geographic sampling with detailed genetic characterization of figs and their pollinators, however, often reveals that multiple pollinators are associated with a single host fig across the range of the fig species (*e.g.*, Michaloud et al., 1996; Haine et al., 2006; Peng et al., 2008; Darwell et al., 2014; Bain et al., 2016). For example, Yu et al. (2019) recovered nine parapatrically distributed pollinator species associated with *Ficus hirta* in South-East Asia. Multiple pollinator species in allopatric and parapatric distributions and associated with the same host suggests pollinator wasps speciate in allopatry before potentially coming back into sympatry. Souto-Vilarós et al. (2019) examined six species pairs of figs and wasps found in New Guinea, and found an increased rate of population genetic divergence with geographic distance among the wasps relative to their host figs. They suggest pollinators have an increased speciation rate, with populations of wasps diverging in isolation faster than their hosts, effectively de-coupling cospeciation processes in the two mutualist taxa (Machado et al., 2005; Peng et al., 2008; Compton et al., 2009; Van Noort et al., 2013; Satler et al., 2019).

Despite the ability of pollinating wasps to disperse many kilometers (Nason et al., 1998; Ahmed et al., 2009), broad geographic sampling suggests that multiple distinct pollinator species—in allopatric and parapatric distributions—characterize host fig species across their ranges. Molecular data suggests a previously underappreciated prevalence of host switching in Neotropical strangler figs, as well as figs in general (Jackson et al., 2008; Cruaud et al., 2012; Yang et al., 2015; Satler et al., 2019). We suspect that combinations of pre- and postzygotic barriers contribute to the reproductive isolation of pollinator wasps (Molbo et al., 2003, 2004; Satler et al., 2019). Specifically, the relative inability of hybrids to detect, locate, and enter a receptive fig, or to their inability to successfully pollinate and oviposit and have offspring successfully develop in a receptive fig, or the effects of hybrid developmental incompatibilities (*e.g.*, hybrid diploid male offspring) would all select against hybrids and for stronger reproductive isolation. When pollinator species experience secondary contact, we suspect these barriers contribute to less fit hybrids and reinforces species boundaries (Nosil et al., 2003).

Consistent with the findings that pollinator species are often well-delimited molecularly, and that F1 hybrids have only rarely been found in the most closely related Panamanian fig wasp species, we expect this to be true among relatively distantly related species (Molbo et al., 2003, 2004; Satler et al., 2019). We suggest that the still poorly understood processes of pollinator speciation and extinction play a more important role than hybridization and introgression in shaping evolutionary dynamics of fig pollinating wasps and their host associations in space and time. Future biogeographical, phylogenetic, and population genetic studies coupled with targeted experimental studies are needed to determine the relative importance of different mechanisms in preventing interspecific mating, hybrid production, and introgression in pollinating fig wasps.

## Conclusions

Given the reproductive biology of fig wasps, multiple events are important for maintaining species boundaries. Before there can be opportunity for hybridization, foundresses from different fig pollinator species need to locate, enter, and oviposit in the same fruit, with successful development of offspring to reproductive age. Although we focused on a Panamanian fig community with an evolutionary history of host switching, and sampled pollinators from hosts that averaged more than one foundress per fruit, we detect no signal of hybridization and introgression among the wasps. The lack of hybridization and introgression indicate that the Panamanian fig pollinators are cohesive evolutionary units. More generally, our results suggest that processes other than interspecific gene flow have been important to the evolution of these fig pollinating wasps.

## Acknowledgements

We thank Brant Faircloth and the Faircloth lab for generously providing time, space, and resources for learning sequence capture approaches for generating UCE data. We thank members of the Heath lab and Nason lab for discussion and comments regarding the manuscript. Funding was provided by the National Science Foundation (DEB-1556853) to JDN, TAH, and EAH, and by the Smithsonian Scholarly Studies Grant to EAH. Computational resources were provided by ResearchIT and the College of Liberal Arts and Sciences at Iowa State University.

## Data Accessibility

Raw sequence data will be available from the NCBI Sequence Read Archive (SRA) under BioProject ID: ### (###). All data sets and custom scripts will be made available on Dryad.

## Author Contributions

JDS, TAH, EAH, and JDN designed the study, AGZ, EAH, and CAM collected the fig wasp samples, JDS generated and processed the sequence data, JDS conducted all analyses, JDS, EAH, and JDN wrote the paper, and all authors contributed to revised versions of the manuscript and approved of the final version.

---

**Figure S1:**
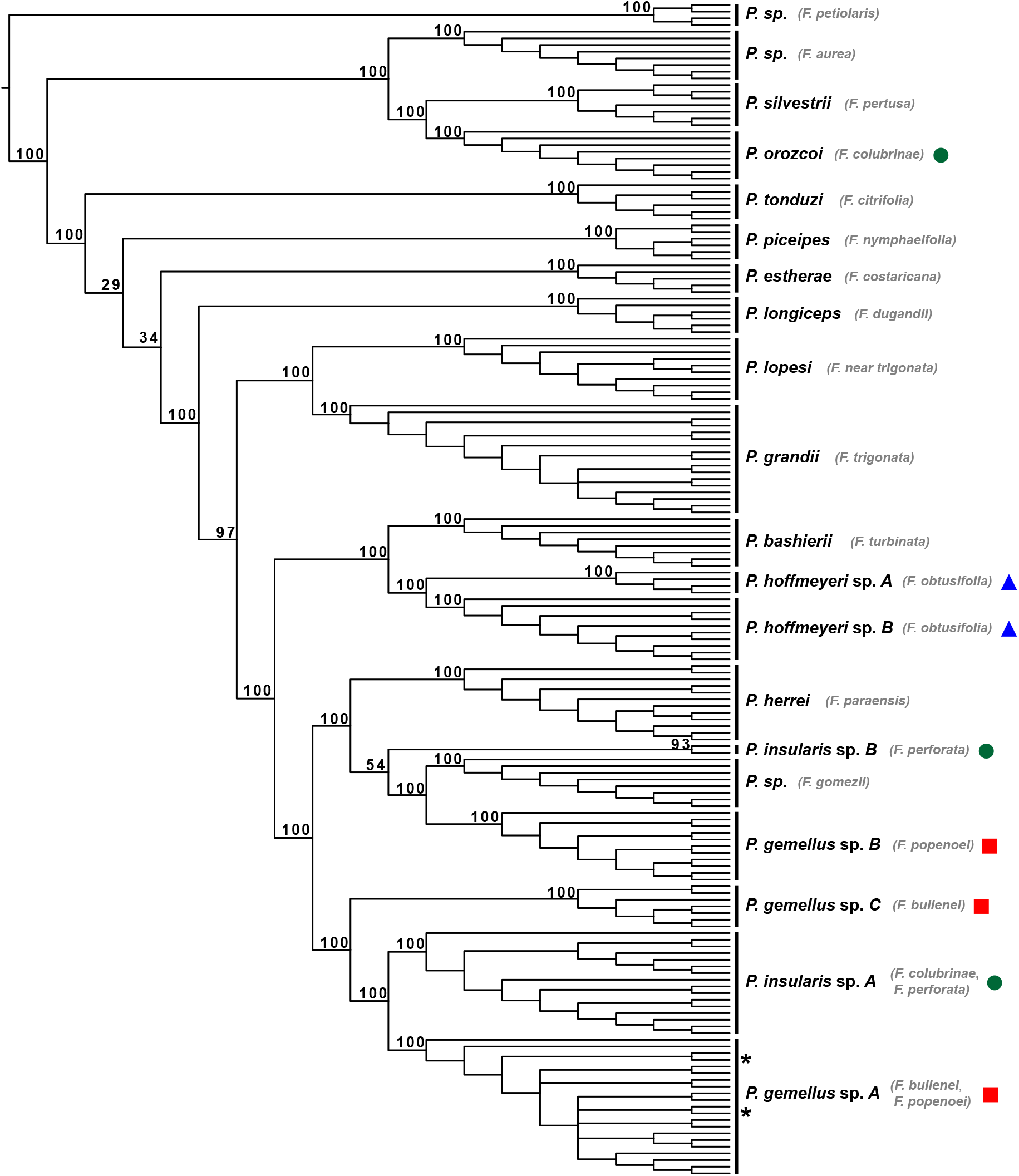
Lineage tree analysis with SVDquartets (SVDQ_*LT*_) where each individual is treated as a tip in the tree. Nodal support reflecting bootstrap values are shown for species and interspecific nodes. Host fig species are displayed next to their associated wasp species. Co-occurring pollinators are denoted by a circle (green), square (red), or triangle (blue). Two individuals (represented by asterisks) of *Pegoscapus gemellus* sp. *A* (hosts: *Ficus bullenei*/*Ficus popenoei*) were sampled from *Ficus dugandii*, which is not its normal host. The undescribed pollinator associated with *Ficus petiolaris* was used to root the tree.

**Figure S2:**
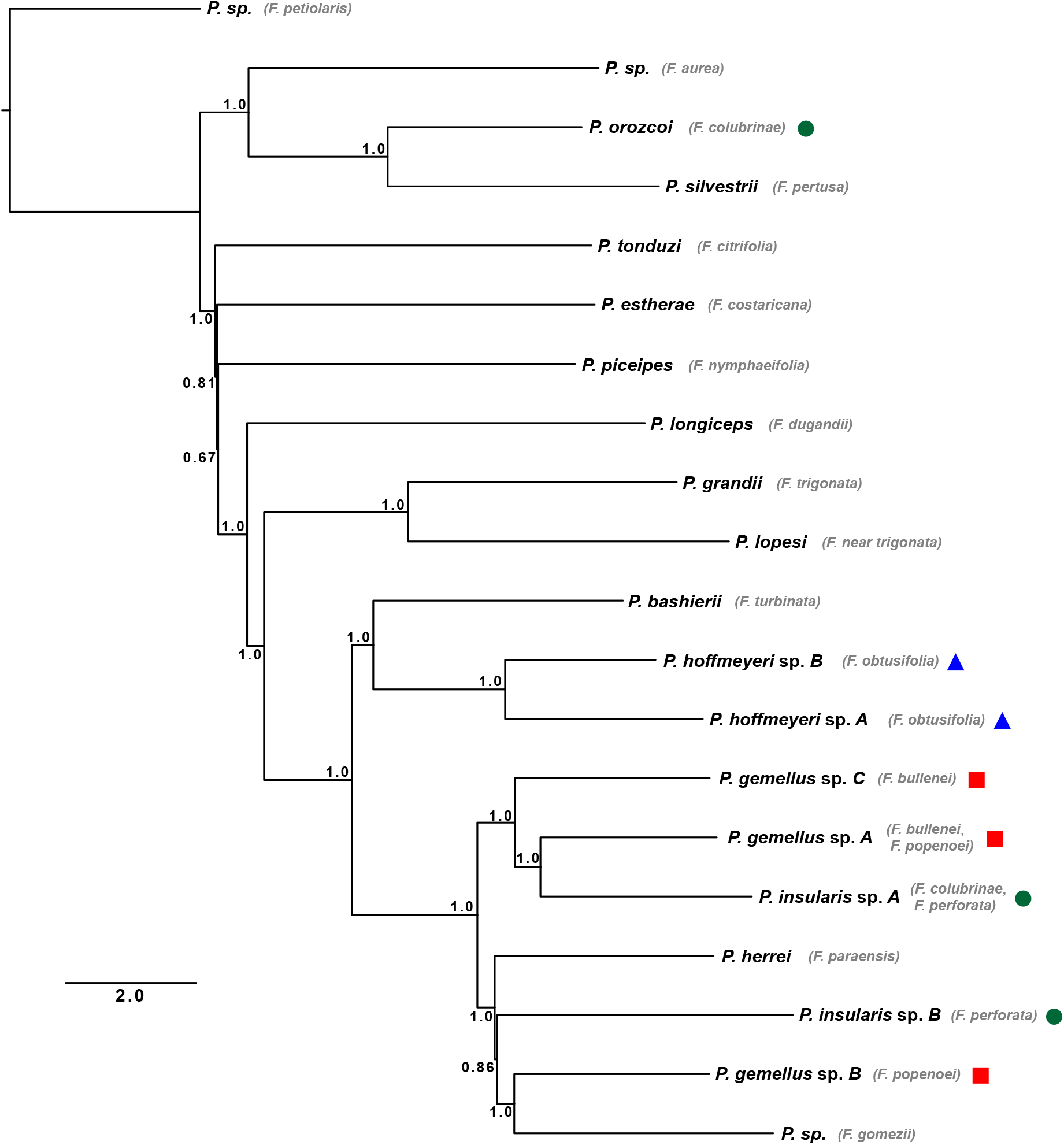
Species tree analysis with ASTRAL-III. Nodal support values represent local posterior probabilities. Host fig species are displayed next to their associated wasp species. Branch lengths are in coalescent units. Co-occurring pollinators are denoted by a circle (green), square (red), or triangle (blue). The undescribed pollinator associated with *Ficus petiolaris* was used to root the tree.

**Figure S3:**
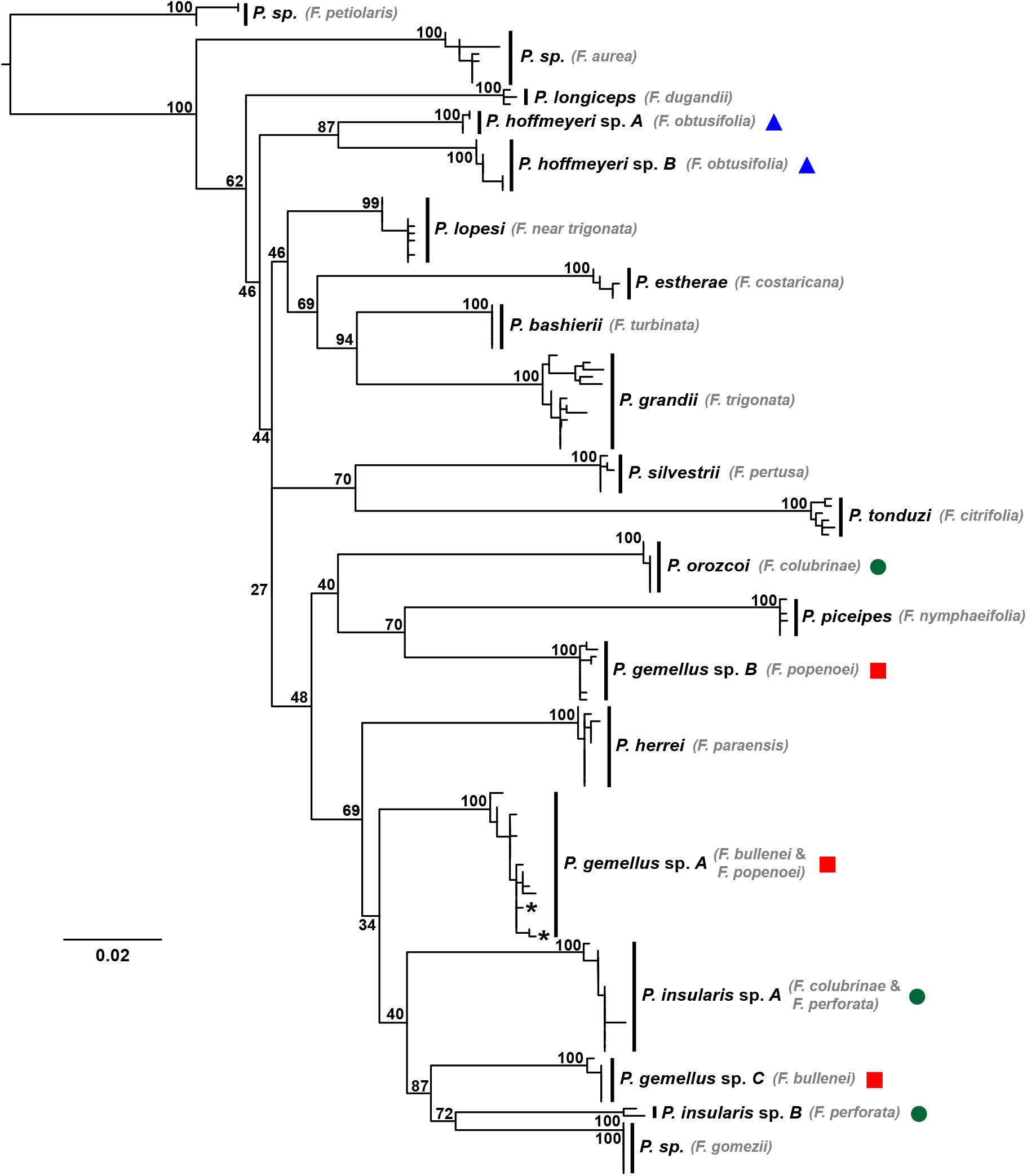
Maximum likelihood gene tree representing relationships among mitochondrial haplotypes. Nodal support reflecting bootstrap values are shown for species and interspecific nodes. Host fig species are displayed next to their associated wasp species. Co-occurring pollinators are denoted by a circle (green), square (red), or triangle (blue). Two individuals of *Pegoscapus gemellus* sp. *A* (hosts: *Ficus bullenei*/*Ficus popenoei*) sampled from *Ficus dugandii* are denoted by asterisks. The undescribed pollinator associated with *Ficus petiolaris* was used to root the tree.

**Figure S4:**
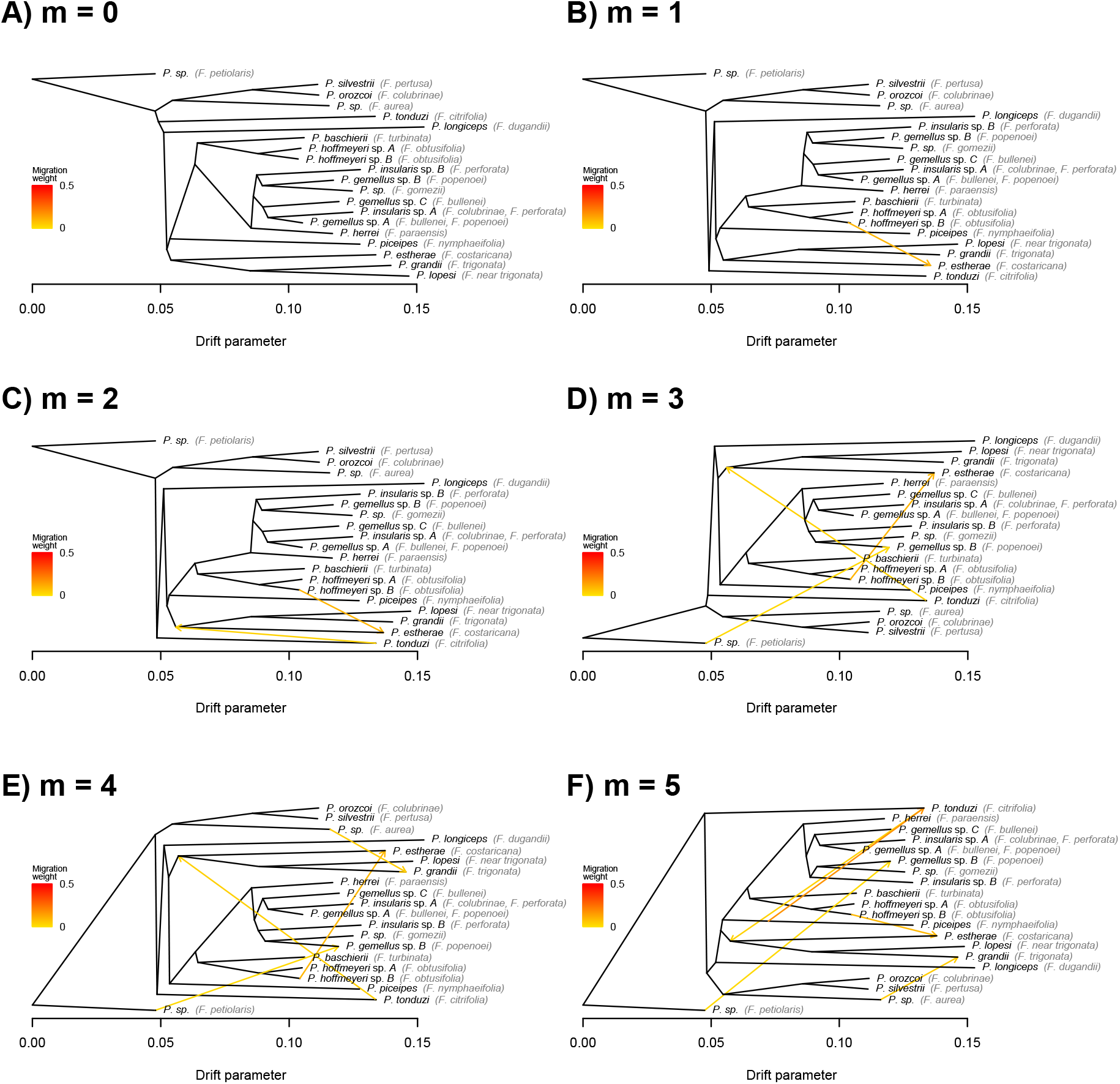
TreeMix results. Results show population graphs without admixture (A), and with between 1 and 5 admixture (m) events (B–F). A model without hybridization (A) is best supported by the data (Table 3). The undescribed pollinator associated with *Ficus petiolaris* was used to root the tree.

